# CRISPR based programmable RNA editing in primary neurons

**DOI:** 10.1101/2023.03.10.532141

**Authors:** Karthick Ravichandran, Tulika Khargonkar, Sarbani Samaddar, Sourav Banerjee

## Abstract

Investigating the RNA regulation landscape primarily relies on our understanding of how RNA-protein interactions are governed in various cell types, including neurons. Analysis of RNA-protein interactions in physiological environments warrants the development of new tools that rely on RNA manipulation. Recently, A CRISPR-based RNA-editing tool (dCas13b-ADAR2_DD_) was developed to mitigate disease associated point mutations in cell lines. Here, we have explored the targeted sequence editing potential of the tool (dCas13b-ADAR2_DD_ system) by adapting it to manipulate RNA function with an aim to visualize RNA editing in primary hippocampal neurons. This is a two-component system that includes a programmable guide RNA (gRNA) complementary to the target RNA, and a catalytically dead version of the Cas13b enzyme fused to ADAR. The RNA editing protocol outlined in this manuscript relies on using the gRNA-dependent targeting of dCas13b-ADAR fusion protein to the mutant form of mDendra transcript. We first abrogated the fluorescence of Dendra2 by introducing a nonsense mutation that precludes the formation of the functional protein. To visualize the efficacy of the RNA editing in neurons, we used the dCas13b-ADAR2_DD_ system to edit specific nucleotides within the Dendra2 mRNA to restore the amino acid codes critical for Dendra2 fluorescence. This method therefore lays the foundation to future studies on the dynamicity of activity-induced RNA-protein interactions in neurons and can be extended to manipulate the endogenous RNome in diverse neuronal subtypes. Furthermore, this methodology will enable investigators to visualize the spatial and temporal resolution of RNA-protein interactions without altering the genomes *via* conventional methods.

**Highlights:**

- CRISPR-Cas13 based application that enables site specific A-to-I in target RNAs directed by gRNA; optimized in neurons.
- Enables the temporal mapping of developmentally relevant RNAs and their cis-interacting RNA binding proteins.

## Background

RNA-based regulatory mechanisms have emerged as key determinants of developmental decisions and cellular metabolism associated with neurons (G. Bassell and Singer, 1997; Doyle and Kiebler, 2011; Kiebler and Bassell, 2006; Liu-Yesucevitz et al., 2011; Martin and Ephrussi, 2009). Activities of both coding and non-coding transcripts rely on their interactions with RNA binding proteins (RBPs) (G. J. Bassell and Kelic, 2004; Darnell, 2006; Shi et al., 2017). These RBPs form Ribonucleoprotein (RNP) complexes that determine diverse RNA metabolism including stability of the transcript, translation, transport, splicing and RNA modifications (Fernandez-Moya et al., 2014; Ramanathan et al., 2019). Current understanding of these RNA metabolic processes stem from disrupting RNA-protein interactions by conventional methods, such as site-directed mutagenesis of RBP binding sites within the target transcript or the inhibition of RBP binding to target transcripts by anti-sense oligonucleotide (Shukla et al., 2020; Taliaferro et al., 2016). Most of the current interaction studies in neurons are structure-based interactions between RNA and their RBP counterparts like Staufen and FMRP (Heraud-Farlow and Kiebler, 2014; Melko and Bardoni, 2010; Mitsumori et al., 2017) However, all of these studies involve the overexpression of exogenous target RNA elements. Secondly, there exist very few reports on the mechanisms underlying cis-acting RNA-RBP interaction (Das et al., 2019). We therefore need unconventional tools to visualize the intricate details of RNA metabolism to an unprecedented resolution.

The discovery of RNA-targeting CRISPR-Cas13 provided a programmable platform for RNA editing and enabled us to visualize RNA metabolism at single transcript resolution (Abudayyeh et al., 2016, 2017; Shmakov et al., 2015). RNA-guided RNA-targeting CRISPR-Cas effector Cas13a from *Leptotrichia wadei* (LwaCas13a), was used for the targeting and knockdown of specific transcripts (Abudayyeh et al., 2017). Later on, Cas13b from *Prevotella sp*. was used for enhanced efficiency of targeting (Cox et al., 2017). The properties of Cas13b enabled the design of a programmable RNA editing system by fusing the catalytically dead version of Cas13b (dCas13b) to ADAR2_DD_; the deaminase domain of ADAR, (<u>a</u>denosine <u>d</u>eaminase <u>a</u>cting on <u>R</u>NA) (Abudayyeh et al., 2017; Cox et al., 2017). ADAR2 is a deaminase that enables RNA editing via the hydrolytic deamination of adenosine to inosine that functionally mimic guanosine during translation (Nishikura, 2010). The RNA guided targeting of dCas13-ADAR2_DD_ to a specific transcript has shown to efficiently edit the RNA in diverse cell lines. However, the application of this system for efficient RNA editing in primary cells, particularly neurons, has not been demonstrated to date.

Primary hippocampal neurons provide a tractable model to study RNA transport and metabolism during different phases of neuronal development. Given their uniquely polar nature, translation at subcellular locations far from the neuronal soma relies on the transport and recruitment of RNAs to translation machineries present in these locales. In order to understand the phenomenology of RNA transport and RNA-RBP association in typical and atypical conditions, it is necessary to elucidate RNA transport and behavior at the resolution of single molecules. Hence we opted to use primary hippocampal neurons as our model system. To demonstrate the efficacy of dCas13b-ADAR2_DD_ –mediated RNA editing system in primary neurons, we have rescued a non-sense mutation of the photoconvertible protein Dendra2; the rescued fluorescence of which serves as a readout of RNA editing. We initially introduced a nonsense mutation in Dendra2; converting the Tryptophan (Trp) codon “UGG” to the translational stop codon “UAG”. This mutation eliminated Dendra2 fluorescence due to premature termination; resulting in no fluorescence detection in primary neurons. Thereafter, we rescued the nonsense mutation in Dendra2 by targeted RNA editing using dCas13-ADAR2_DD._ Successful RNA editing was determined by the recovered Dendra2 fluorescence.

Recently developed RNA editing techniques involve the recruitment of endogenous ADARs in a guide RNA dependent manner. These methods, owing to longer guide RNA lengths, might promote ADAR hyperactivity leading to non-specific adenosine de-aminations within the target RNA (Wahlstedt and Öhman, 2011). Contrarily, the dCas13-ADAR2_DD_ is augmented through rational modifications within ADAR. These increase specificity of edits within the target RNA sequence, while causing less transcriptome-wide off-target edits overall. Moreover, pathological conditions where ADAR activity is subject to variability, through mutations or altered expression levels (Slotkin and Nishikura, 2013), may in turn make editing events unpredictable and irreproducible. The dCas13-ADAR2_DD_ circumvents this by editing in a controlled, dosage dependent manner. We envisage that this programmable RNA editing can be further extended to study state-dependent diverse RNA metabolic processes in subcellular locations within neurons.

## Materials and Reagents

### Biological Materials

1. Timed pregnant CD1 mice (E17) maintained at Animal House, National Brain Research Center
2. Stellar^™^ Competent Cells – Clontech #636763
3. pDendra2-C – Clontech #632546
4. pC0052-REPAIR non-targeting guide clone into pC0043 – Addgene plasmid #103868
5. pC0055-CMV-dPspCas13b-GS-ADAR2DD(E488Q/T375G)-delta-984-1090 – Addgene plasmid #103871

### Reagents

1. Lipofectamine-2000TM – Invitrogen #11668019
2. Trypsin 2.5% – Gibco #15090046
3. Poly(L-Lysine-Hydrobromide) – Merck #P2636
4. Neurobasal medium – Gibco #21103049
5. B27 supplements – Gibco #17504001
6. HEPES – Gibco #11344041
7. Hanks’ Balanced Salts – Sigma-Aldrich, #H2387
8. Sodium bicarbonate – Sigma Life Science #S5761-500G
9. Penicillin-Streptomycin – Gibco #15140-122
10. Minimal Essential Media – Gibco # 11095-072
11. Cytosine β-D-Arabinofuranoside, Hydrochloride – Sigma-Aldrich #C6645
12. 100X Pyruvate – Gibco #11360-070
13. Horse Serum – Gibco #26050-070
14. QIAprep Spin Miniprep Kit - Qiagen #27106
15. EndoFree Plasmid Maxi Kit - Qiagen #12362
16. Potassium acetate – Merck #61870305001730
17. Potassium hydroxide – Sigma-Aldrich #221473-500G
18. Magnesium Acetate Tetra-hydrate – Sigma Life Science #M5661-250G
19. T4 DNA Ligase - New England Biolabs #M0202S
20. T4 Polynucleotide Kinase - New England Biolabs #M0201S
21. ATP 10mM - New England Biolabs #P0756S
22. Nitric Acid 69% - Merck #1.93006.2521
23. Luria-Bertani Broth (Miller) Microbiology – Merck #1.10285.0500
24. Ampicillin Sodium Salt – Sigma-Aldrich #A9518-25G
25. UltraPure™ DNase/RNase-Free Distilled Water - Invitrogen #10977023

### Solutions

1. 5x Annealing Buffer
2. HEPES-KOH
3. Dissection Media
4. Poly-L-Lysine Solution
5. Neurobasal-B27
6. Glial MEM

### Recipes

1. 5x Annealing Buffer

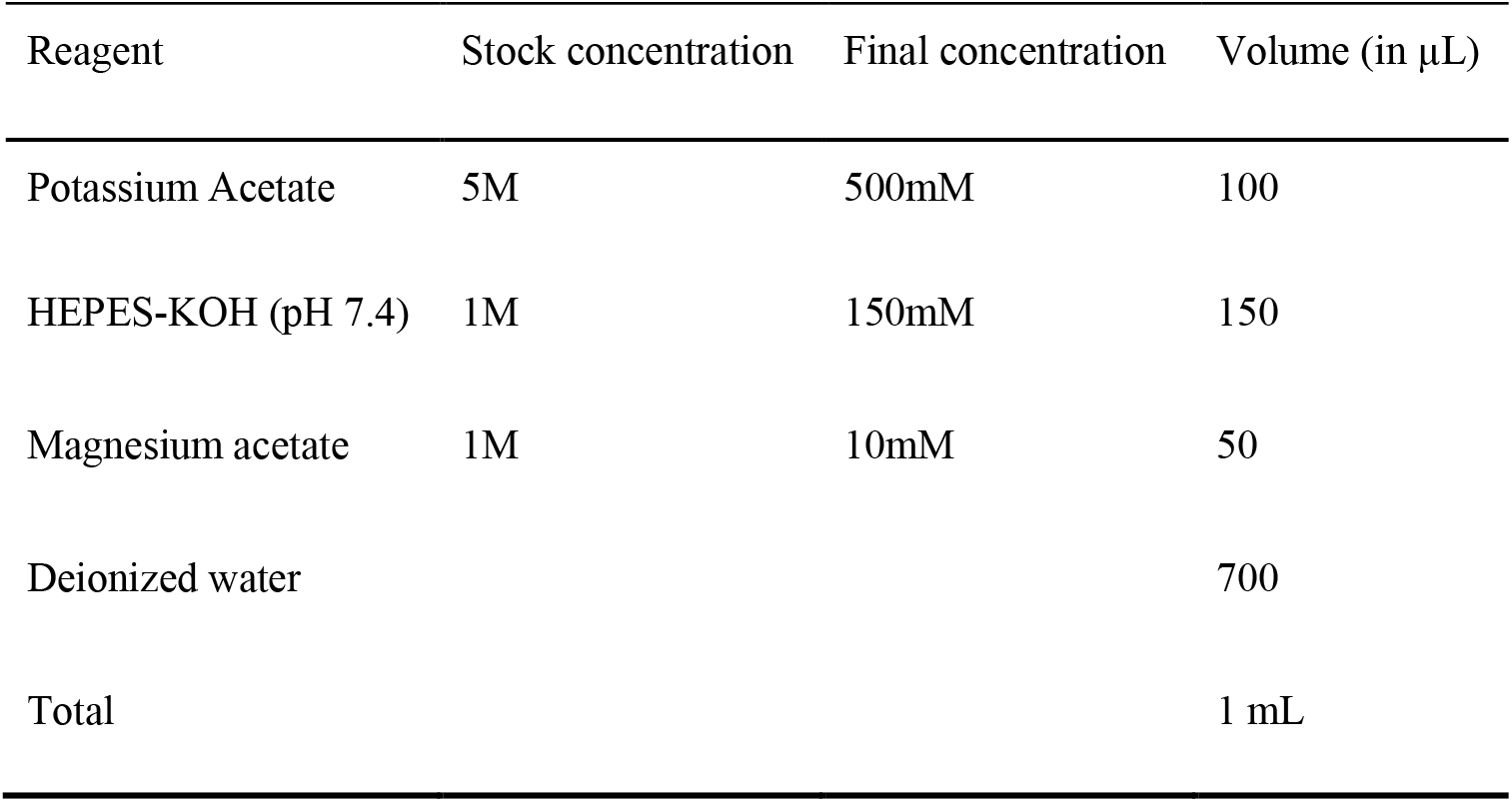
2. HEPES-KOH Dissolve 23.8g in 80mL of MilliQ water and adjust the pH 7.4 using 10N KOH. Adjust the final volume to 100mL using MilliQ. Filter using 0.22μm filter and store at 4°C.
3. Dissection Media

4. Poly-L-Lysine Solution

| Reagent | Amount |
| --- | --- |
| Hank's Balanced Salt | 9.5 g |
| Sodium Bicarbonate | 0.35 g |
| HEPES-KOH pH7.4 (1M) | 20 mL |
| Penicillin-Streptomycin | 20 mL |
| MilliQ | Make up to 1L |
| Total | 1000 mL |

Filter with a 0.22µm filter. Store at 4°C.

5. Neurobasal-B27 Media

Filter with a 0.22µm filter. Store at 4°C. 4. Poly-L-Lysine Solution
| Reagent | Final concentration | Amount |
| --- | --- | --- |
| Poly-L-Lysine | 0.1mg/mL | 1g |
|  | 20 mM | 2 mL |
| H<sub>2</sub>O | n/a | 95 mL |
| Total | n/a | 100 mL |

6. Glial MEM

5. Neurobasal-B27 Media
| Reagent | Amount |
| --- | --- |
| Neurobasal | 500 mL |
| B-27 Supplement | 10 mL |
| L-Glutamine (200mM) | 1.25 mL |
| Penicillin-Streptomycin | 1 mL |
| Total | 512.5 mL |

Filter with a 0.22µm filter. Store at 4°C.

6. Glial MEM
| Reagent | Amount |
| --- | --- |
| MEM | 425 mL |
| 20% Glucose | 15 mL |
| 100X Pyruvate | 5 mL |
| Penicillin-Streptomycin | 5 mL |
| Horse Serum | 50 mL |
| Total | 500 mL |

### Equipment

1. Forma CO_2_ Incubator – ThermoScientific #381
2. Water bath circulator – Neolab
3. Heating Block – Neolab
4. Milli-Q® Direct Water Purification System – Merck #ZR0Q008WW
5. Incubator Shaker (New Brunswick Excella E25 or equivalent)
6. Dissection Microscope – Zeiss Stemi 2000-C Stereo Microscope #455053-0000-000 or equivalent
7. Nunc^™^ Cell-Culture Treated Multidishes (or equivalent) – ThermoScientific #150628
8. Cover glasses – Blue Star

### Procedure

#### A. Preparation of primary mouse hippocampal neuronal culture

Primary hippocampal culture from mouse neurons was prepared as per the protocol mentioned in (Kaech and Banker, 2006) with modifications. Here, the hippocampus is dissected from brains of embryonic day 18 mouse pups. A single cell suspension is generated through trypsinization and trituration, and seeded onto poly-L-lysine coated cover glasses in neurobasal media-containing 12-well plates.

The culture is stable for a period of 28 days. However, the quality of the culture degrades with time, hence the suitable time window for editing is before 8 *days in vitro*. Protocol can be adapted to cell culture plates and dishes available at hand.

##### 1. Prepare culture dishes as indicated below

a. Place 18mm cover glasses on ceramic racks and immerse them in nitric acid (69%) overnight.
b. Remove the ceramic racks and place in autoclaved MilliQ water for 30 minutes.
c. Repeat the rinse with fresh MilliQ water three more times.
d. Drain the water. Enwrap the ceramic racks in aluminum foil.
e. Bake in a hot air oven at 150°C overnight or a minimum of 16 hours. Do not unwrap the oven baked cover glasses outside the laminar chamber.

##### 2. Coat the culture dishes as indicated below

a. Transfer sterile 12-well plates and the ceramic rack containing cover glasses for culture into the Laminar flow chamber.
b. Carefully place the cover glasses into sterile 12-well plates using blunt-end forceps.
c. Sterilize the laminar chamber along with the culture dishes under UV radiation for a minimum of 1 hour.
d. Coat the 18mm cover glasses with refrigerated poly-L-lysine hydrobromide solution.
e. Cover the culture dishes with aluminum foil and leave them in the chamber overnight (before the day of dissection) *The poly-L-lysine hydrobromide solution can be recovered and stored aseptically. This can be reused; however, the number of washing steps has to be reduced accordingly*.
f. Aspirate the poly-L-lysine hydrobromide into a sterile container.
g. Wash the culture dishes with autoclaved MilliQ water at room temperature for 10 minutes.
h. Repeat the wash twice (for fresh poly-L-lysine).
i. Add Glial MEM into the culture dishes and place them in the incubator.

##### 3. Dissect the mouse hippocampus

a. Prepare the following dissection tools:
b. Euthanize time pregnant CD1 mouse (E17-18) in a carbon dioxide chamber. Confirm death by checking for movement on toe and tail pinch.
c. Place the mouse facing upwards and sterilize its peritoneal region with 70% Ethanol (v/v)
d. Using sterile toothed forceps and Mayo scissors, make an incision perpendicular to the midline in the peritoneum. Extend the incision through the midline until the thorax (Figure 1).

**Figure 1:**
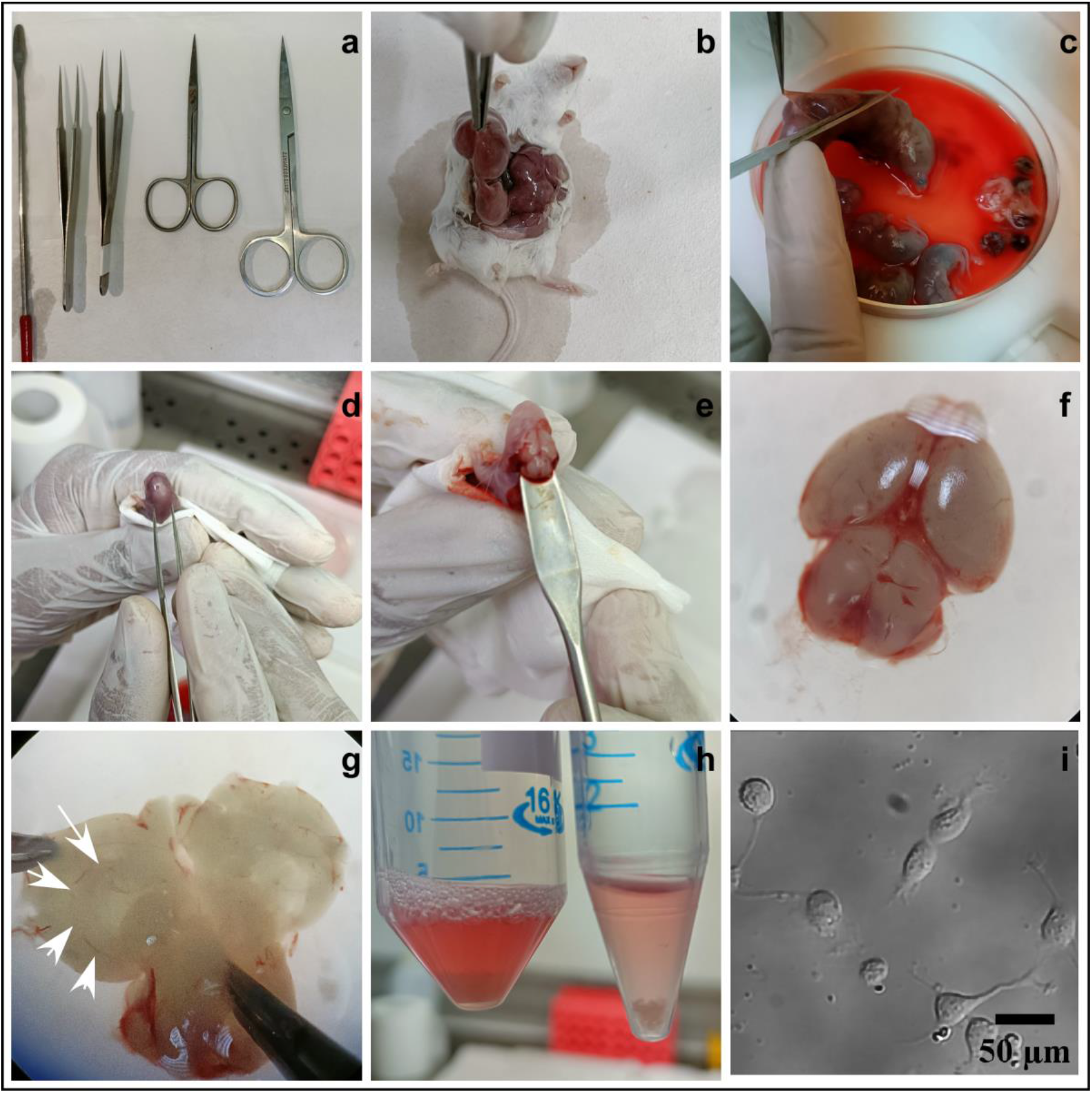
Demonstration of mouse brain removal and hippocampal dissection. **A**. Surgical instruments B. Extraction of pups from uterus C. Extraction of pups from amniotic sac D. Removal of overlapping skin and skull E. Removal of brain from pup F. Brain under dissection microscope G. Separation of hemispheres; hippocampus visible H. Collected hippocampal fractions immersed in Dissection Media I. Bright field image of plated neurons.
Remove the uterus and place it in ice cold dissection media in a 90mm petri dish. *The extraction of the pups can be performed outside the laminar chamber. The pups should be maintained in ice cold dissection media. Further microdissection of the hippocampus from the pups should be carried out in a controlled sterile laminar chamber*.
Transfer the uterus and surgical apparatus for microdissection into a sterile laminar chamber.
Extract the pups using forceps and Iris scissor.
Hold the pup in one hand with an open palm positioned between the thumb and forefinger. With fine forceps, remove the skin covering the head.
Using a microdissection scissor, make an incision at the bottom of the skull and extend it through the midline till the nose. Make a lateral cut and carefully pull off the skull without damaging the brain.
Scoop the brain and place it on the petri dish containing dissection medium.
Using fine forceps separate the hemispheres and pull the hemisphere laterally such that the sagittal plane becomes visible under the dissection microscope.
Hippocampus can be identified as a “C” shaped structure. Using fine forceps, dissect the bilateral hippocampi from the brain. Perform the dissection for all the pups.
Transfer the hippocampi to a fresh 15mL centrifuge tube.

##### 4. Preparation of hippocampal neurons for plating

a. Make up the volume of the centrifuge tube with dissection media. To this, add 20µL of 2.5% Trypsin per pup (Time pregnant mice typically have 6-11 pups). Incubate the tube at 37°C in a water bath for 15 minutes.
b. Remove the supernatant without disturbing the hippocampi. Make up the volume to 5mL with warm dissection media, to neutralize trypsin activity.
c. Incubate at room temperature for 10 minutes. Repeat the washes with warm dissection media twice.
d. Decant the dissection media from the tube. Add 2mL of pre-warmed Glial MEM and dissociate the hippocampal tissue by repeated pipetting using 1mL micropipette.
e. Triturate by passing the tissue through a fire-polished Pasteur pipette for 10 to 15 passes, until the chunk disappears. *The Pasteur pipette is subjected to a fire to narrow its diameter. However, an excessively narrow diameter will cause cell damage and affect viability*.
f. Make up the cell suspension volume to 3mL with Glial MEM.
g. Enumerate cell count using a hemocytometer. Cells can now be seeded as per requirement.

### Maintenance of hippocampal culture

1. Add cytosine arabinoside or *AraC* to a final concentration of 1-5µM on *day in vitro*/DIV 3. *Addition of AraC prevents the growth of glial cells in the culture*
2. Replace half of the existing media in the culture plates with fresh Neurobasal-B27 media. Post *DIV 7*, add glial conditioned Neurobasal-B27 media in addition to the fresh media (100μL per mL).
3. Periodically, visually examine the culture dishes for turbidity, indicative of bacterial or fungal contamination. pH changes can be observed on change in media color, indicative of biological contamination or cell death.

#### B. Targeted editing of RNA

Here we provide general instructions for the design of guide RNA to be cloned into the expression vector. Once successfully cloned, dCas13b-ADAR2_DD_ and guide RNA constructs are to be transfected into the prepared neuron culture.

Editing efficiency will depend on the chosen transfection method, health of neurons at the time of transfection, and specificity of guide RNA design.

##### 1. Design of guide RNA against target RNA

a. The guide RNA will consist of two parts—a 30 nucleotide-long direct repeat sequence for dCas13b construct recognition, and a 50 nucleotide-long spacer specific for target sequence.
b. The direct repeat sequence, as suggested by (Cox et al., 2017). – 5’ – GUUGUGGAAGGUCCAGUUUUGAGGGGCUAUUACAAC – 3’ The direct repeat will be at the 3’ end of the guide RNA. Hence this sequence should directly follow the spacer sequence.
c. A poly-T track of 8 nucleotides should be included following the direct repeat sequence, which terminates RNA polymerase activity.
d. The target adenosine to be deaminated should be preferentially mispaired with a cytosine within the spacer sequence in the guide RNA.
e. Mismatch distance is defined as the number of nucleotides between the target adenosine and the start of the direct repeat sequence. It should be empirically determined depending on experimental conditions. *A mismatch distance of 16 nucleotides resulted in highest editing rates in our experiments*.
f. For cloning the guide RNA into the expression vector, restriction site adaptors should be appended at the ends of the guide RNA sequence.
g. The guide RNA should be designed in accordance with basic primer design principles.
  - Base composition should be 40-60% (G+C).
  - Melting temperatures or T_m_ should range between 55-80°C.
  - T_m_ values should be within 5 degrees of each other.
  - There should be no sequence stretches with self-complementarity.

##### 2. Cloning of guide RNA into expression vector

a. pC0052-REPAIR non-targeting guide clone into pC0043 – is obtained from Addgene (Addgene plasmid #103868).
b. Resuspend the guide RNA oligonucleotides to the same molar concentration in nuclease-free water.
c. Anneal the guide RNA oligonucleotides – mix equimolar amounts of the guide RNA oligonucleotides in nuclease-free water and annealing buffer. In a thermocycler, perform annealing with the following parameters:

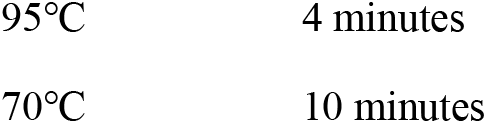
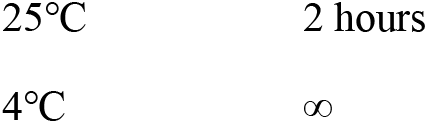
d. Perform polynucleotide kinase treatment to phosphorylate the guide RNA – mix 5µg of guide RNA, with appropriate volumes of T4 kinase 10x buffer, 10µM ATP, T4 kinase and nuclease free water. In a thermocycler, perform PNK treatment with the following parameters:

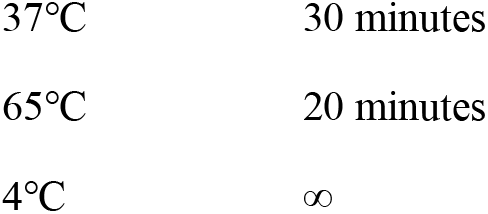
e. Clone the guide RNA into pC0052 directly downstream of the U6 promoter. Multiple cloning sites can be used to insert the guide RNA in a directional manner. *In this case, pc0052 was double digested using AgeI and Acc65I*. *The gRNA was designed such that the AgeI restriction site was located on the 5’ end, while Acc65I restriction site was located on the 3’ end*.
f. Digest pc0052 using chosen restriction enzymes.
g. Ligate the guide RNA and pc0052. Prepare a ligation mixture in nuclease-free water, 10x T4 buffer, and T4 ligase. Incubate at room temperature for 10 minutes, followed by incubation at 65°C in a water bath *or* a heat block for 10 minutes. *Suggested molar insert to vector ratio is 7:1. NEBioCalculator® can be used for ligation calculations*.
h. Transform the plasmid into Stellar Competent cells. Allow them to grow on an ampicillin-containing agar plate overnight.
i. Pick isolated colonies and inoculate them in LB Broth overnight. *Inoculate in appropriate volume of LB Broth depending on the nature of the DNA isolation kit*.
j. Perform DNA extraction using commercially available DNA isolation kits. *Use of an endotoxin-free maxi-prep kit is recommended*.
k. Verify the sequence of the product using available primers.

##### 3. Transfection delivery of constructs into cultured neurons

*The editing transfection mix will contain*

a. *A plasmid encoding dCas13b-ADAR2*_*DD*_ *construct*,
b. *A plasmid encoding the guide RNA, and*
c. *Based on use case, a plasmid encoding exogenous target RNA*.

*Amount of DNA to be transfected will vary depending on transfection conditions, method used, and number of neurons to be transfected, and needs to be empirically determined. dCas13b-ADAR2*_*DD*_ *construct to guide RNA construct ratio should be 1:2*.

a. Warm neurobasal media at 37°C for an hour prior to transfection.
b. Aspirate media from neuron dishes. Supplement it with B-27–this is hereafter referred to as conditioned media.
c. Replace with the pre-warmed neurobasal media.
d. Prepare the transfection mix in the laminar chamber.
e. Add 100μL of neurobasal media to an eppendorf tube labelled A. To this, add DNA corresponding to the transfection conditions.
f. In a separate eppenddorf tube labelled B, add 100 μL of neurobasal media. Add 1 μL of Lipofectamine-2000 for every 1 μg of DNA to be transfected. Incubate at room temperature for 5 minutes.
g. Mix the contents of tubes A and B to create the transfection mix. Pipette gently and incubate at room temperature for 20 minutes.
h. Add the transfection mix to the neuron dishes drop-wise. Mix by swirling on the bench.
i. Place the dishes in the incubator for 1 hour.
j. Aspirate media from the dishes. Wash with fresh neurobasal media thrice to remove transfection mix. *Swirl and tap the dishes while performing the wash step to ensure dead cells and transfection debris is removed properly. Improper washing may result in additional cell death*.
k. Replace with conditioned media supplemented with B-27.
l. Incubate the neurons for 48 hours post-transfection to allow Cas13b-mediated RNA editing to occur.

## Results

The dCas13-ADAR2_DD_ along with a guide RNA construct were expressed in primary neurons. The guide RNA targets the dCas13-ADAR2_DD_ to the site of nonsense mutation in Dendra2 *via* its sequence complimentary binding on either side of the mutation. The *specificity of edit* was primed by the “cytidine” base flip in the RNA duplex of gRNA-Dendra2 mRNA which led to ADAR–mediated deamination of the “Adenosine” of “UAG” stop codon to “Inosine”. The translation apparatus recognizes this inosine for its equivalent guanosine, and incorporates a Trp amino acid in the polypeptide chain, rescuing premature termination and restoring Dendra2 fluorescence properties (Figure 4).

**Figure 2:**
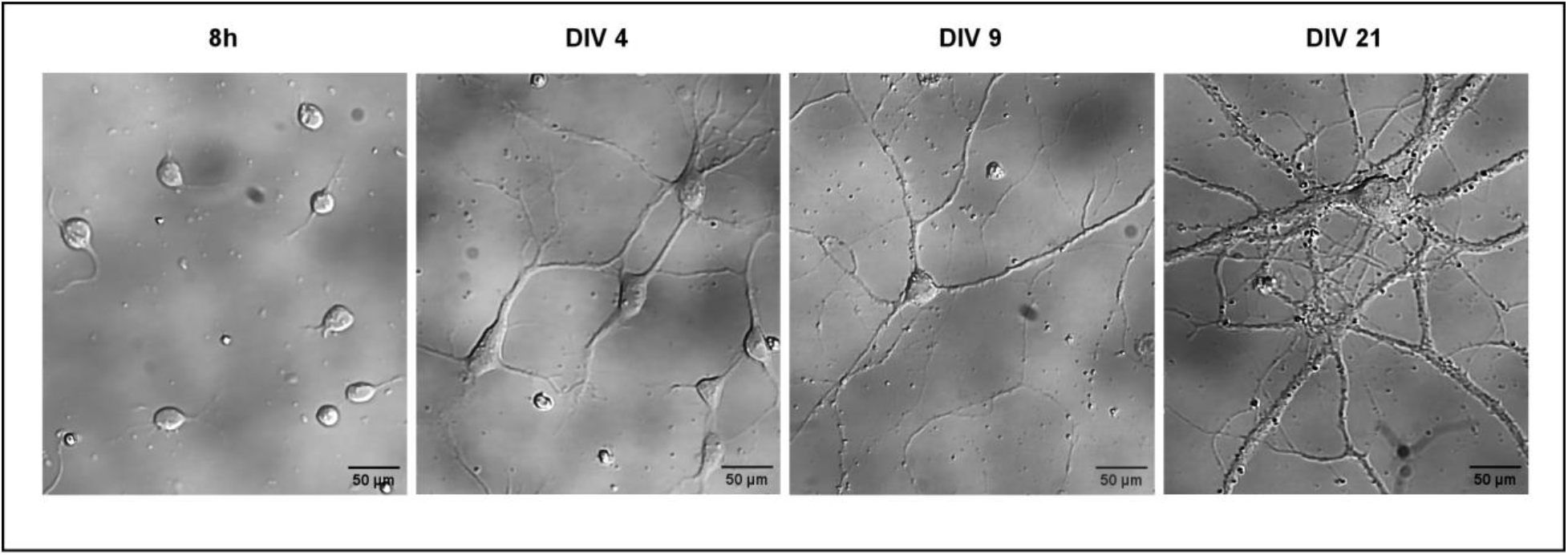
Low density plated primary hippocampal neurons at definite developmental time points. Figure depicts developmental stages of cultured hippocampal neurons *in vitro* at DIV 0, DIV4, DIV 9 and DIV 21.All panels represent DIC images of neurons taken at 40X magnification in a bright-field microscope. Scale bar as indicated.

**Figure 3:**
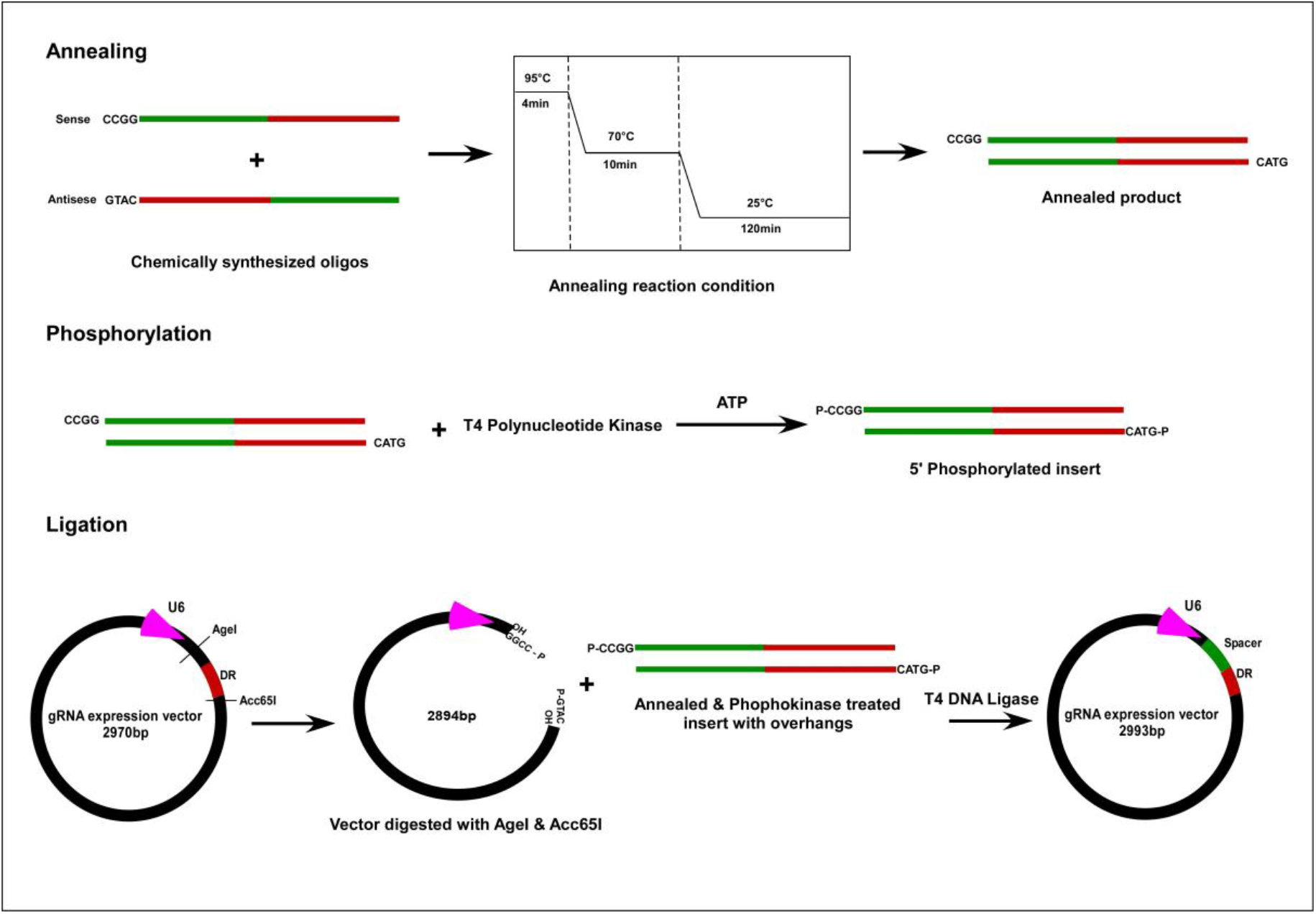
Schematic of guide RNA cloning. Designed inserts (obtained as synthetic DNA oligonucleotides) are annealed, phosphorylated using T4 Polynucleotide Kinase and ligated using T4 DNA Ligase to digested vector.

**Figure 4:**
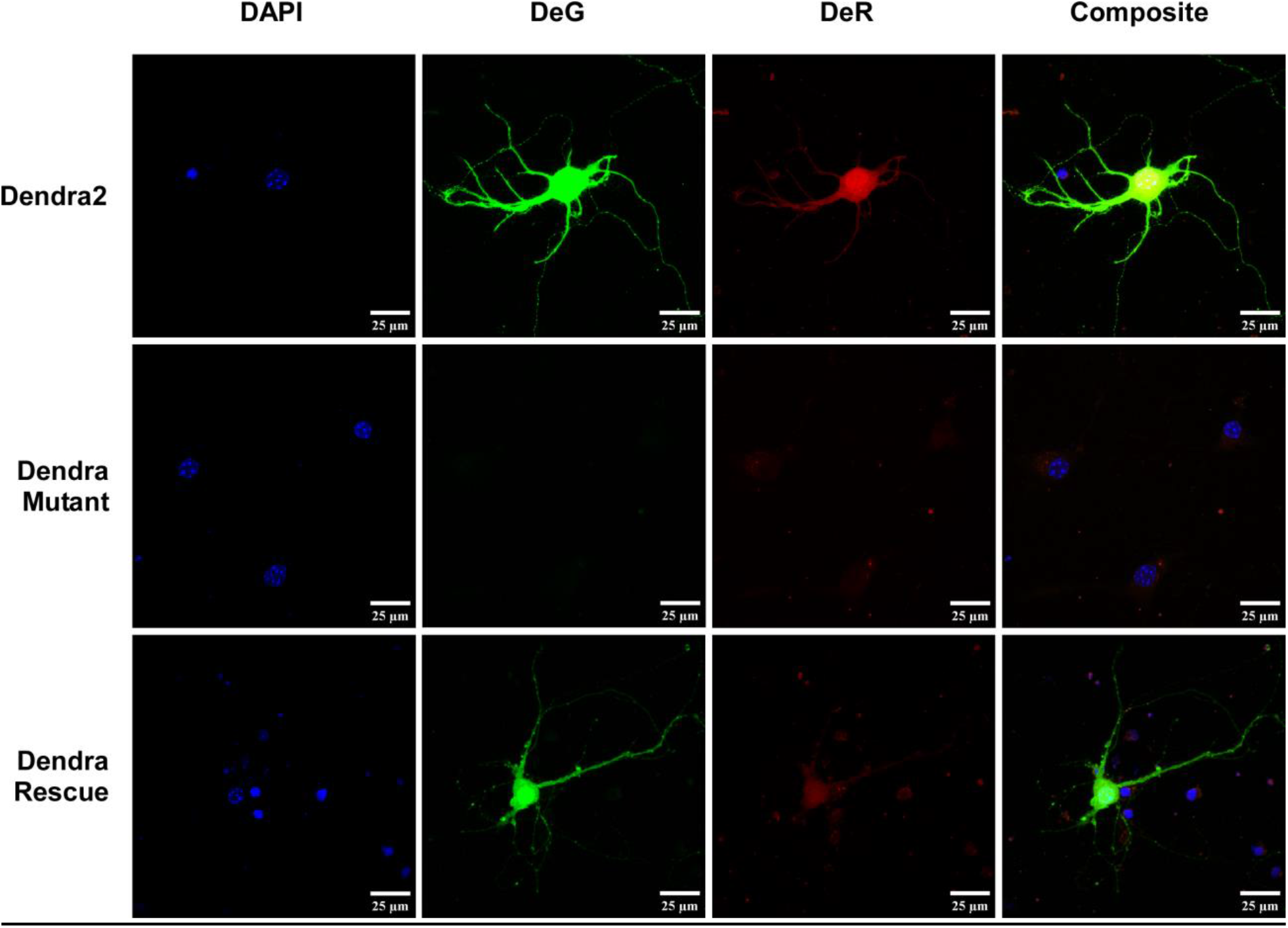
Dendra2 rescue using the dCas13b-ADAR2_DD_ system. Top panel shows native Dendra2 protein fluorescence acquired in both native green (DeG) and red (DeR) channel. Middle panel shows the mutated Dendra2 that constituted no detectable fluorescence in green and red channel. Bottom panel shows the rescue of fluorescence in green and red channel. DAPI is used as nuclear stain. All images were acquired using *Nikon* A1 HD microscope at 100x magnification. Scale bar as indicated in figure.

### Troubleshooting

| Problem | Possible Cause | Solution |
| --- | --- | --- |
| Formation of cell clumps | Insufficient trituration | Triturate with the number of passes suggested in the protocol; Ensure that the Pasteur pipette diameter is sufficiently narrow |
|  | High seeding density | Seed lesser number of cells |
| Cell death post-seeding | Excessive trituration | Triturate with the number of passes suggested in the protocol |
| Cell death post-transfection | Improper washing | Ensure removal of precipitates from the media |
| Low transfection efficiency | Insufficient DNA : lipofectamine complex formation | Empirically optimize the DNA: lipofectamine ratio; Increase DNA : lipofectamine complex formation time |
| Low editing efficiency | Incorrect guide RNA design | Follow the protocol for guide RNA design |

## Discussion

The dCas13b-ADAR2_DD_ system is unique because its dose-dependency is user-regulated. This system allows the manipulation of mRNA transcripts at the single molecule level without affecting either its stability or its secondary structure.

Study of RNAs and cis-interacting protein partners is largely studied with the lens of secondary structure-specific interactions. However some literature makes a case (Singer) for sequence specific involvement of RNAs. Structure and sequence resolution study of these interactions, taken in tandem, might present a clearer picture of their nature.

A growing body of literature affirms the importance of the transcriptome—coding and non-coding alike—in neuron-specific phenomena of synaptic plasticity, neurodevelopment, etc.

## Acknowledgments

This study was supported by Department of Biotechnology, India grant BT/PR31811/GET/119/285/2019. The protocol is adapted from by Cox *et al*. (2019).

## Ethical considerations

All animal handling procedures approved by the IAEC (Institutional Animal Ethics Committee) of NBRC (National Brain Research Centre), India, under the protocol numbers (NBRC/IAEC/2020/167).

The NBRC IAEC is registered with the CPCSEA (Committee for the Purpose of Control and Supervision of Experiments on Animals) (Registration number: 464/GO/ReBi-S/Re-L/01/CPCSEA) the Ministry of Fisheries, Animal Husbandry and Dairying, Government of India.

